# A plasmid-encoded type IIB restriction-modification system in cyanobacteria blocks conjugative gene transfer

**DOI:** 10.1101/2025.10.21.683720

**Authors:** Brenda Pratte, Teresa Thiel

**Affiliations:** Department of Biology, University of Missouri-St. Louis, One University Dr., St. Louis, MO, 63130, USA

## Abstract

*Anabaena* (aka *Trichormus*) *variabilis* ATCC 29413 is a filamentous, heterocyst-forming cyanobacterium with a 6.36 Mb chromosome, four circular plasmids, A (366 kb), B (35.8 kb), C (301 kb), and D (27 kb), and a 37-kb excision element. The *hsdRMS* genes on plasmid D may encode a type IIB restriction-modification system that protects the cells from invasion by foreign DNA. A variant of *A. variabilis*, strain FD, lacking plasmid D, grew with the same generation time as *A. variabilis* ATCC 29413, which stably maintained plasmid D. *Nostoc* sp. PCC 7120 and *Nostoc* sp. M131, which lack plasmid D and *hsdRMS*, grew similarly with a synthetic replicating plasmid, with or without added *hsdRMS* genes. Although the natural plasmid D was very stable in *A. variabilis* ATCC 29413, the synthetic plasmid was easily lost in *Nostoc* sp. PCC 7120 and *Nostoc* sp. M131, with or without *hsdRMS*. Strain FD, lacking plasmid D, and an *A. variabilis hsdRM* deletion mutant were much better hosts for conjugation of a non-replicative, integrative plasmid than wild-type *A. variabilis*. The conjugative nonreplicating plasmid formed circular molecules in strain FD and in the *A. variabilis hsdRM* deletion mutant, allowing for a high percentage of single-recombinant exconjugants. In contrast, the relatively few exconjugants in *A. variabilis* were all double recombinants, suggesting that the restriction-modification system encoded by the genes on plasmid D resulted in only linear molecules that recombined by double crossovers. Expression of the *hsdRMS* genes in strains *Nostoc* sp. PCC 7120 and *Nostoc* sp. M131 drastically reduced conjugation frequency and recombination of a non-replicative plasmid. Together, these data indicate that HsdRMS acts as a defense against the successful transfer of foreign DNA into these cyanobacteria, likely by double-stranded DNA cleavage. Many cyanobacterial strains that are not amenable to gene transfer might have similar restriction-modification systems that lead to cleavage and subsequent degradation of foreign DNA. Although such systems inhibit gene transfer, they may yield a high proportion, although a low number, of gene replacement recombinants.

## Introduction

*Anabaena (*aka *Trichormus) variabilis* ATCC 29413 (hereafter, *A. variabilis*) is a filamentous, heterocyst-forming cyanobacterium. The original deposited genome sequence consists of a 6.36 Mb chromosome, three circular plasmids A (366 kb), B (35.8 kb), and C (301 kb), and a 37-kb linear element (Thiel et al., 2014; GenBank assembly ID GCA_000204075). Resequencing nominally the same strain (Mardanov et al., 2013; Thiel et al., 2014) showed the existence of a fourth plasmid, D (27 kb), that was also seen in physical experiments (Lambert and Carr, 1982; Simon, 1978). The apparent anomaly was resolved when we resequenced our lab strain, confirmed that it lacks plasmid D, and realized that the strain sequenced by JGI that we provided to them was our lab variant strain of *A. variabilis*, called FD (Currier and Wolk, 1979). We also sequenced six other cyanobacterial strains, originally cultured from the water fern *Azolla* (Zimmerman et al., 1989), that are nearly identical to *A. variabilis* ATCC 29413 (Pratte and Thiel, 2021). *A. variabilis* ARAD, V5, 9RC, N2B, FSR, and PNB all have plasmid D, in addition to the other plasmids (although *A. variabilis* FSR lacks part of plasmid A (Pratte and Thiel, 2021). Therefore, the absence of plasmid D in strain FD suggests that its loss was the result of growing the parent strain *A. variabilis* for many generations at 40°C (Currier and Wolk, 1979), which was not known previously.

Plasmid D has *hsdRM* and *hsdS* genes for a putative restriction/modification (RM) system, *parAB* genes for putative plasmid partitioning, and genes for hypothetical proteins (Mardanov et al., 2013). RM systems recognize and cleave foreign DNA with unmethylated bases in the endonuclease recognition sequence (Vasu and Nagaraja, 2013) to protect the cell from invasion by foreign DNA (Arber, 1979). The *hsdRM* and *hsdS* genes were reported to constitute a type I system (Mardanov et al., 2013); however, the genes are similar to those encoding type II restriction enzymes in which a single, composite HsdRM enzyme has both the restriction and modification activities, whereas HsdS is similar to the type I S subunit that specifies the DNA recognition sequence for the RM system. (Marshall and Halford, 2010; Marshall et al., 2007; Smith et al., 2012). *A. variabilis* has two type II restriction endonucleases, AvaI and AvrII (Duyvesteyn et al., 1983; Herrero et al., 1984).

We hypothesize that the HsdRMS system on plasmid D protects cyanobacterial cells from invading foreign DNA, including phages and plasmids. Thus, the lack of *hsdRMS* in strain FD might account for its ability to better support phage infection (Currier and Wolk, 1979) and possibly gene transfer compared to the parent strain, *A. variabilis*. Here, we characterized *hsdRMS* by determining the rate of loss of the plasmid under non-selective conditions and the effect of the addition of the *hsdRMS* genes on growth rate and gene transfer efficiency.

## Results and Discussion

### Bioinformatic analysis of HsdRM and HsdS in plasmid D

The *hsdRM and hsdS* genes on plasmid D in *A. variabilis* were reported to be a type I system, but the evidence, based on the similarity to characterized proteins, is weak (Mardanov et al., 2013). Based on conserved motifs, the predicted proteins for *hsdRM* (locus tag Fig. 1. Plasmid D (NC_021575.1) and conserved motifs in HsdRM and HsdS. Motifs for HsdRM and HsdS in *A. variabilis* were identified using the MOTIF program provided by GenomeNet (https://www.genome.jp/tools/motif/). HTB18_RS00005) and *hsdS* (locus tag HTB18_RS00020) may form a type IIB RM system, in which the HsdR and HsdM domains are fused into one subunit, similar to BcgI (Kong et al., 1994) (Marshall and Halford, 2010; Marshall et al., 2007; Smith et al., 2012). Type IIB systems make a double-stranded break on either side of the recognition site, releasing a short 34-bp DNA fragment for BcgI (Marshall et al., 2007). HsdRM in *A. variabilis* has the PD….(D/E)XK endo motif (amino acids 444-452, **PD**VSSL**DPK**) characteristic of the bipartite catalytic site in type II systems (Kosinski et al., 2005), and the motif order—the Motif I sequence (amino acids 375-386, **V**M**DP**AC**G**S**G**G**FL**), and the Motif IV sequence (amino acids 498-507, **FD**V**IL**A**NPPF**)—typical of a γ-class N6-methyl adenine DNA methyltransferase (Madhusoodanan and Rao, 2010) (Malone et al., 1995) (**Fig. 1**). The separate type I S-like subunit has two putative target recognition domains, TRD1 and TRD2, potentially providing a bipartite target recognition sequence suggesting that the enzyme cleaves outside of the recognition sequence (**Fig. 1**). All of this is consistent with HsdRMS functioning as a type IIB restriction-modification system. In addition to domains for endonuclease and methylase activities, HsdRM has a motif of unknown function that has some similarity to chromosome segregation proteins. Further characterization of the cleavage sites will be necessary to determine whether it is a type IIB system.

**Fig. 1.**
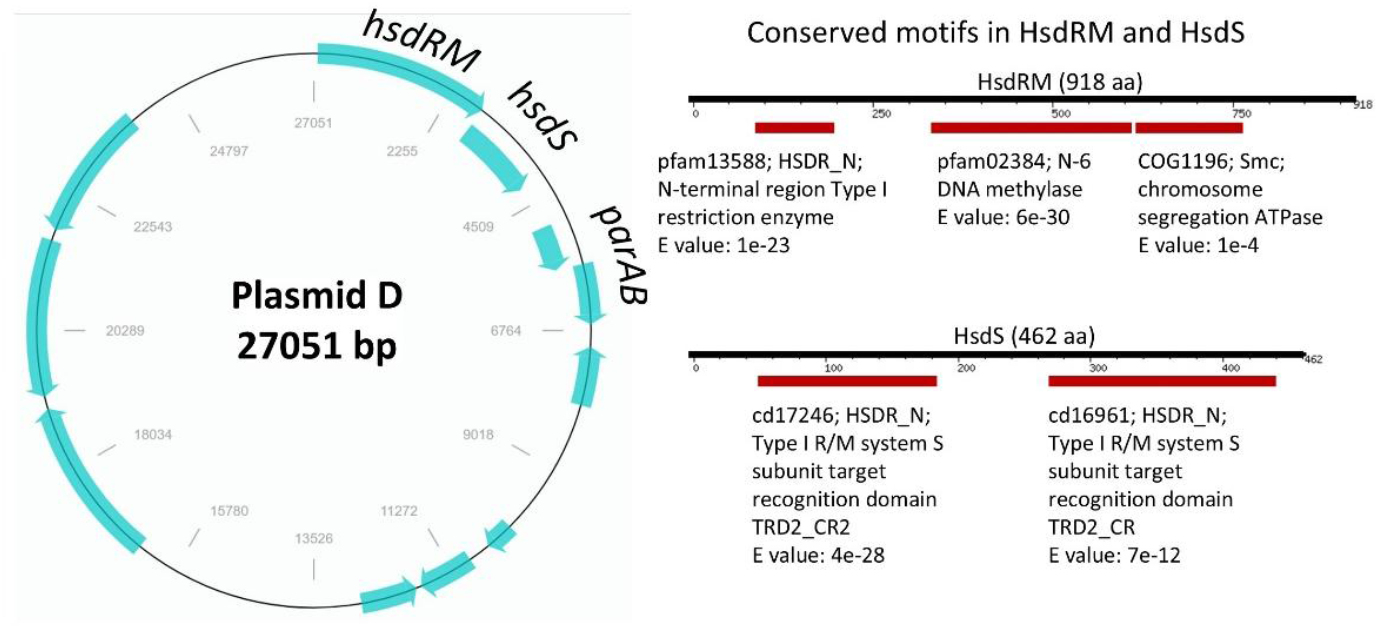
Plasmid D (NC_021575.1) and conserved motifs in HsdRM and HsdS. Motifs for HsdRM and HsdS in *A. variabilis* were identified using the MOTIF program provided by GenomeNet (https://www.genome.jp/tools/motif/).

### Plasmid copy number and hsdRMS expression

Since only *A. variabilis* and its close relatives have plasmid D with the *hsdRMS* genes, we cloned these genes into the replicating plasmid pRL57, which is based on the *Nostoc* plasmid pDU1 (Reaston, 1982; Wolk et al., 1984). The resulting plasmid, pBP1207, was transferred to *Nostoc* sp. M131 and *Nostoc* sp. PCC 7120 by conjugation. Plasmid D in *A. variabilis* had an average copy number of 1.3 compared to the chromosome, similar to the previously reported value of one (Mardanov et al., 2013) and much lower than the pDU1-based plasmid pBP1207, which had a copy number of about 12 in *Nostoc* sp. PCC 7120 and *Nostoc* sp. M131. The parent plasmid pRL57 had about the same copy number in *Nostoc* sp. M131, but an anomalously high copy number of about 33 in *Nostoc* sp. PCC 7120 (see **Fig. 4**). Plasmid copy number for pDU1-based plasmids has been reported to vary enormously between about 0.5–1800 copies per chromosome depending on the host strain, plasmid insert, and growth conditions (Wolk et al., 2007; Yang et al., 2013). Expression of *hsdRMS* in the *Nostoc* strains with pBP1207 was more than 50-fold higher than the native, low-copy number plasmid D in *A. variabilis*, 5-fold higher than one would expect based solely on the difference in gene dosage (**Fig. 2**).

**Fig. 2.**
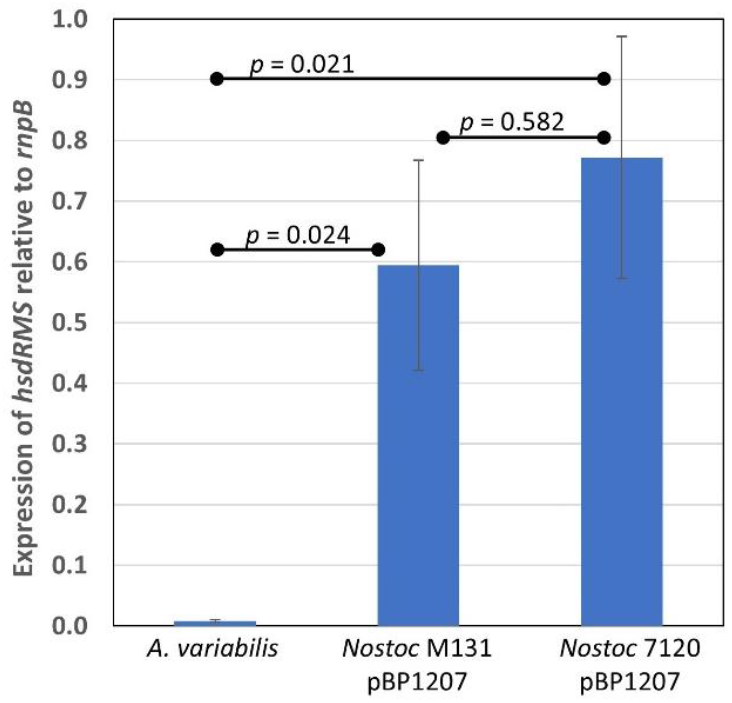
Expression of *hsdRMS*. The level of expression of *hsdRMS* relative to *rnpB* was determined by RT-qPCR for the strains indicated in the graph. *p*-values are indicated between the values for the two strains that were compared.

### Effect of hsdRMS on growth and loss of plasmids

Strain FD, lacking plasmid D with the *hsdRMS* genes, and the parent strain *A. variabilis* (with plasmid D) had generation times of about 4 h during exponential growth in dilute cultures that minimize self-shading and light limitation (**Fig. 3**). In contrast, *Nostoc* sp. PCC 7120 and *Nostoc* sp. M131, with either pRL57 (no *hsdRMS*) or pBP1207 (with *hsdRMS*), had much longer generation times than *A. variabilis* and FD. The presence of the *hsdRMS* genes in *A. variabilis* on plasmid D and in *Nostoc* sp. M131 and *Nostoc* sp. PCC 7120 on plasmid BP1207 did not affect the growth of these strains compared to strains lacking the *hsdRMS* genes. In the absence of antibiotic selection, pBP1207 was lost slowly in both *Nostoc* sp. M131 and *Nostoc* sp. PCC 7120, with a loss of about 50– 75% of the copies over nine generations. However, similar plasmid loss also occurred in strains with the control plasmid pRL57, suggesting that the presence of the *hsdRMS* genes was not associated with plasmid loss. This plasmid loss was surprising, as these plasmids are based on the endogenous *Nostoc* sp. PCC 7524 plasmid, pDU1 (Reaston, 1982). Thus, it is likely that genes in pDU1 that are lost in pRL57 and pBP1207 contribute to plasmid defense and stability. Further, the artificial plasmids are likely maintained in high copy number only by the presence of antibiotics; hence, removal of antibiotic selection leads to loss of the plasmid.

**Fig. 3.**
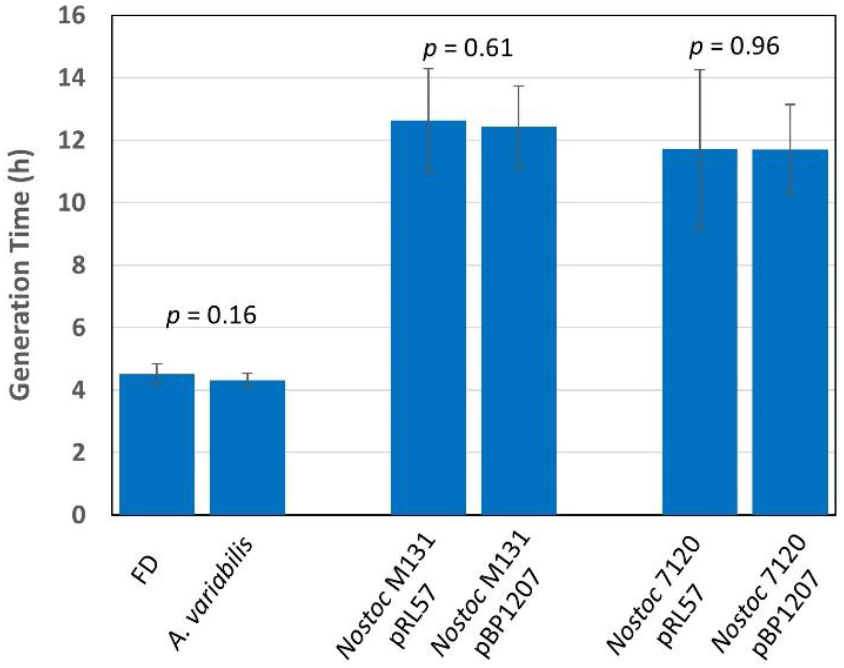
Growth rates of strains with or without *hsdRMS*. The generation times of *Nostoc* sp. M131 or *Nostoc* sp. PCC 7120 with plasmids pRL57 (no *hsdRMS*) or pBP1207 (with *hsdRMS*) and FD (without plasmid D) or *A. variabilis* (with plasmid D) were determined as described in the Methods. *p*-values are indicated above the two values that were compared.

We had no selection for plasmid D in *A. variabilis*; therefore, we could not attempt to cure plasmid D from *A. variabilis* by removing selection. However, strain FD, lacking plasmid D, was originally obtained by growing *A. variabilis* at 40°C (Currier and Wolk, 1979), suggesting that this relatively high temperature might lead to loss of plasmid D. We grew *A. variabilis* for nine generations at 40 °C but found no decrease in plasmid D copy number over that time (**Fig. 4**). Subsequently, we grew the strain for over 60 generations at 40°C but saw no change in the average plasmid copy number, which would likely reflect the combination of cells with no plasmid and those with a normal copy number. If we assume that we would detect a loss of 50%, then the rate of loss of plasmid was < 1.1% per generation [(1-p)^60^ > 0.5]. Although *Nostoc* sp. M131 and *Nostoc* sp. PCC 7120 lost copies of pBP1207 over only a few generations, that loss cannot be attributed to the *hsdRMS* genes since pRL57, lacking *hsdRMS*, was also lost. The maintenance of the natural plasmid D in *A. variabilis* may confer advantages to the cell that pBP1207 does not in *Nostoc* sp. M131 or *Nostoc* sp. PCC 7120. However, the natural plasmid D may be purely parasitic with defense systems against loss, whereas the artificial plasmids, pRL57 and pBP1207, have no such defense.

**Fig. 4.**
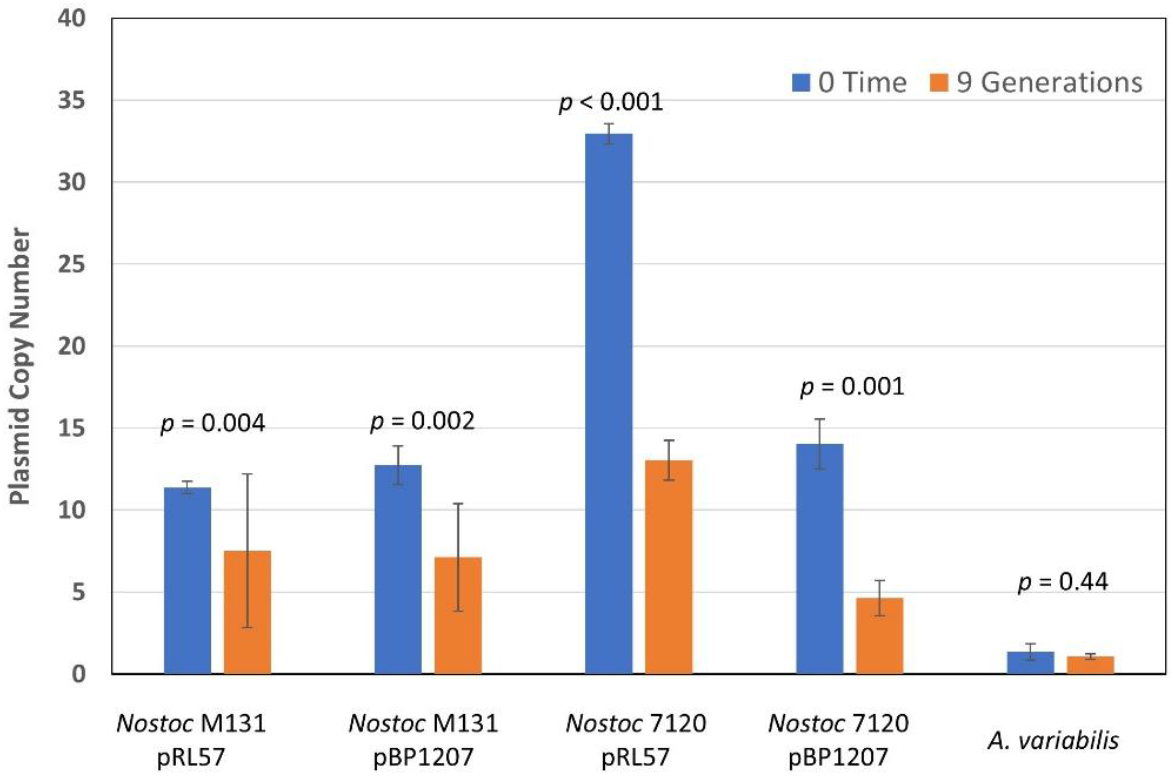
Plasmid copy number and plasmid loss. The plasmid copy number relative to the chromosome (measured by *rnpB*) was determined by qPCR for cells prior to (0 time) and 9 generations after either removal of antibiotic selection (for strains with plasmids pRL57 or pBP1207) or growth at 40 °C (for plasmid D in *A. variabilis*). *p*-values are indicated above the two values that were compared.

### Conjugation efficiency and single versus double recombination

The FD variant of *A. variabilis* was originally reported to support phage N1 infection better than the parent strain (Courier and Wolk, 1979), and which could represents a more general phenomenon: a decreased defense against foreign DNA. To test this, we determined the efficiency of conjugation for *A. variabilis* and FD by conjugating plasmid pBP201, containing a 2.5-kb *modAE* (molybdate transport genes) (Zahalak et al., 2004) region from the *A. variabilis* chromosome, cloned in a mobilizable vector that cannot replicate in *A. variabilis*. The plasmid has an Nm^R^ gene in the middle of *modE* and an Em^R^ gene in the vector for selection of recombination of the plasmid with the chromosome (Nm^R^), and screening for double *modAE* recombinants lacking the vector portion of the plasmid (Em^S^). As molybdate is only required for nitrogenase and nitrate reductase (Mouncey et al., 1995), growth with ammonia is not affected by the loss of molybdate uptake.

The plasmid pBP201 transferred well and integrated using the *modAE* region into the chromosome in strain FD, which lacks *hsdRMS*. However, it conjugated poorly to *A. variabilis*, which has *hsdRMS*, with an apparent efficiency of only about 5–10% relative to FD (**Table 1**). Although plasmid pBP201 can integrate into the *modAE* region of the chromosome by either single recombination (Nm^R^Em^R^ colonies) or double recombination (Nm^R^Em^S^ colonies), single recombinants are statistically the most common. This was true for exconjugants of FD (87–90% single recombinants), but the very few exconjugants in *A. variabilis* were all double recombinants. In *A. variabilis* strain BP1309, in which *hsdRM* was deleted in plasmid D, conjugation restored the high frequency of *modAE* single recombinants; however, the number of exconjugants was far fewer than in FD, suggesting that other genes in plasmid D may contribute to the low frequency of gene transfer (**Table 1**). The presence of the *hsdRM*S genes in *Nostoc* sp. PCC 7120 and *Nostoc* sp. M131 abolished exconjugants (except for one colony for *Nostoc* sp. PCC 7120). In the absence of HsdRMS, the plasmid was successfully integrated by single or double recombination in *Nostoc* sp. PCC 7120 and *Nostoc* sp. M131; however, the frequency of double recombinants was low compared to single recombinants (**Table 2**).

**Table 1.**
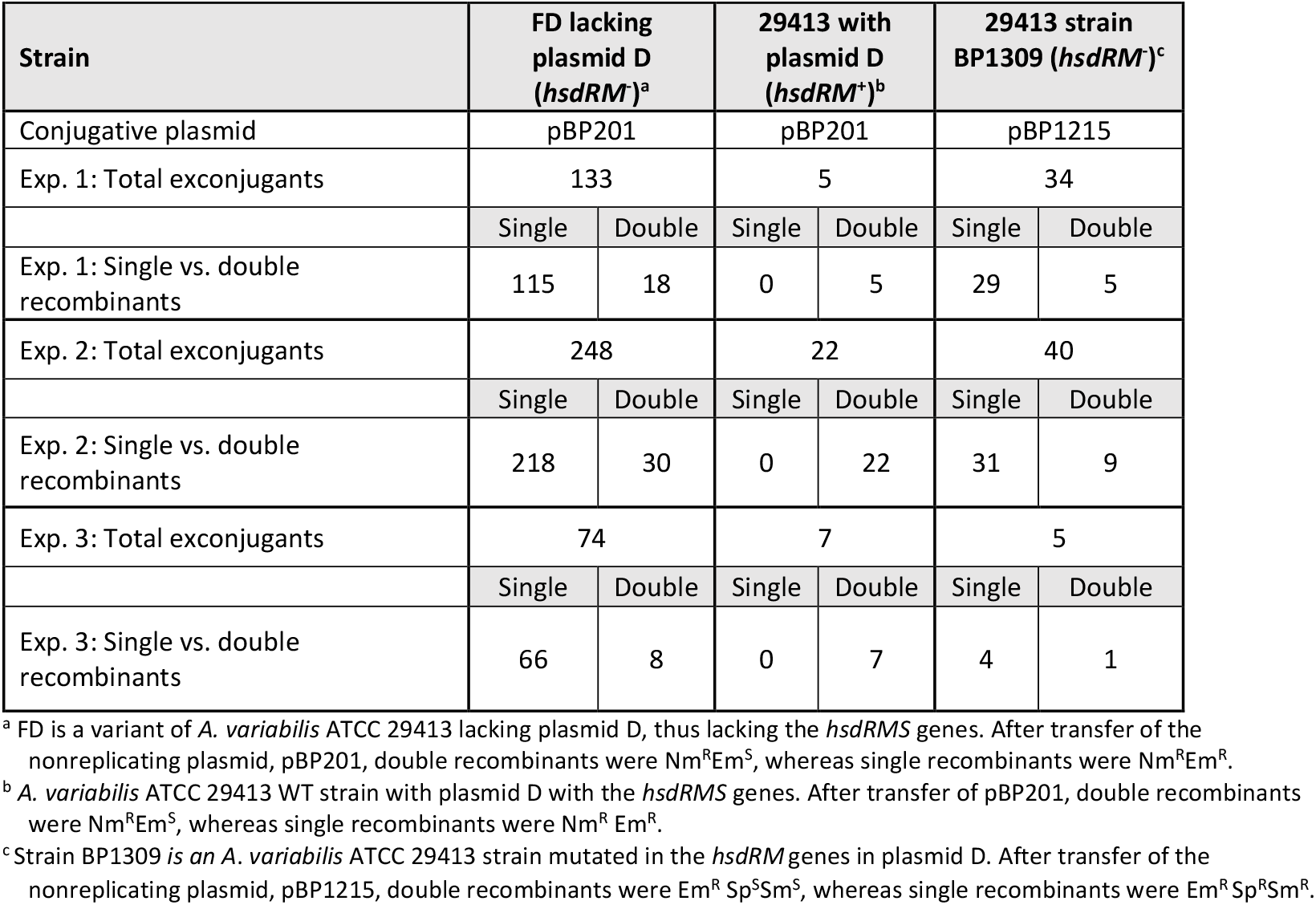
Conjugation of a non-replicative plasmid into *A. variabilis* and FD strains ±*hsdRMS* and numbers of single vs. double recombinants.

**Table 2.**
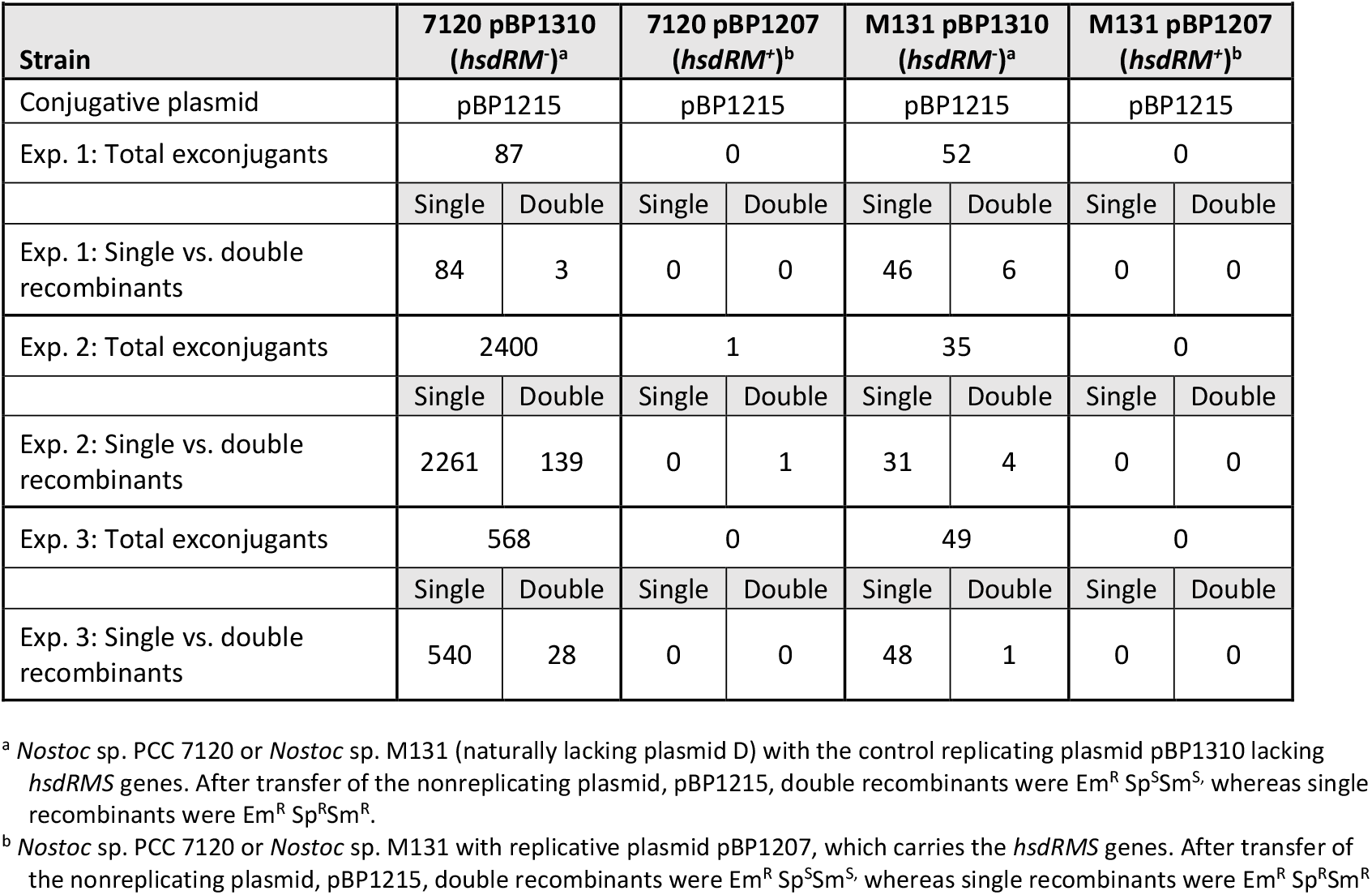
Conjugation of a non-replicative plasmid into *Nostoc* 7120 and M131 strains ±*hsdRMS* and numbers of single vs. double recombinants.

Circular DNA can yield either single or double recombinants, whereas linear DNA can yield only double recombinants. A single recombination event between the circular chromosome and the linear plasmid would linearize the chromosome, which could be lethal. One possible explanation for the loss of single recombinants is that HsdRMS mediates cleavage of the conjugative plasmid after it enters the cell. This would result in linear plasmids or plasmid fragments that could integrate only by double recombination, yielding a low frequency of exconjugants. However, it may not be only HsdRMS that results in double recombinants, since FD, lacking this nuclease, produced 10–15% double recombinants. These double recombinants may result from double recombination with a circular plasmid, or another endonuclease may linearize the incoming plasmid. The negative effect of restriction endonucleases on conjugation efficiency of non-integrating plasmids in cyanobacteria has been demonstrated (Elhai et al., 1997; Wolk et al., 1984). Further, other restriction sites in the plasmid could yield a linear plasmid in FD. As the *modAE* region has no AvaI or AvrII sites, that region would be immune to cleavage by these restriction enzymes. There are four AvaI and two AvrII sites in the vector region of pBP201, and seven AvaI sites and no AvrII sites in the vector region of pBP1512. However, the helper conjugation plasmid in *Escherichia coli*, pRL528, encodes an AvaI methylase that should protect the transferred plasmid from Ava I cleavage in FD (Elhai and Wolk, 1988a). While HsdRMS drastically decreased conjugation efficiency, it also provides selection for double recombinants. Thus, bacterial strains with these or similar restriction enzymes could be useful in experiments in which only double recombinants leading to gene replacement are desired.

## Summary and Conclusions

*A. variabilis* plasmid D is not essential for growth. A variant of *A. variabilis*, strain FD, lacks plasmid D but grows with the same generation time as *A. variabilis*. Plasmid D is very stable in *A. variabilis*, and several strains nearly identical to *A. variabilis* ATCC 29413 (*A. variabilis* ARAD, V5, 9RC, N2B, FSR, and PNB) have plasmid D. The *hsdRMS* genes on plasmid D appear to encode a type IIB RM system that protects the cells from invasion by foreign DNA. Strain FD, lacking plasmid D, was a much better host for conjugation than *A. variabilis*, and conjugative plasmids formed circular molecules in FD, allowing for a high percentage of single-recombinant exconjugants. In contrast, linear plasmid molecules in *A. variabilis*, likely resulting from cleavage of the transferred plasmid, yielded only double recombinants. Together, these data indicate that HsdRMS acts as a defense against the successful transfer of foreign DNA into these cyanobacteria, likely by double-stranded DNA cleavage of the foreign DNA at unknown sites. Many cyanobacterial strains that are not amenable to gene transfer might have similar RM systems that lead to cleavage and subsequent degradation of foreign DNA. An RM system in such a strain could be inactivated by the insertion of an antibiotic resistance gene into the RM gene by double crossover recombination after conjugation with a plasmid carrying the appropriate gene construct. However, the RM system could also provide selection for double recombinants.

## Methods

### Strains and growth conditions

Strains and derivative strains of *A. variabilis* 29413, *A. variabilis* variant FD, *Nostoc* sp. PCC 7120, and *Nostoc* sp. M131 were maintained on BG-11 agar medium (Rippka et al., 1979) supplemented, when appropriate, with 50 μg ml^−1^ neomycin sulfate (Nm), 3 μg ml^−1^ spectinomycin and streptomycin (SpSm) or 5 μg ml^−1^ erythromycin (Em). Strains were grown photoautotrophically in liquid cultures in an eight-fold dilution of AA medium (AA/8)(Allen and Arnon, 1955) supplemented with 5 mM NH_4_Cl and 10 mM TES, pH 7.2, at 30 °C, with illumination at 50–80 μeinsteins m^−2^ s^−1^. Antibiotics were added to liquid cultures as follows: Nm (5 μg ml^−1^), SpSm (0.5 μg ml^−1^), and/or Em (1 μg ml^−1^).

Cultures used to determine growth rate were initially grown to an OD_720_ of 0.2 in 125-ml flasks containing 50-ml AA/8 supplemented with 5 mM NH_4_Cl and 10 mM TES, pH 7.2, and, when appropriate, 5 µg ml^-1^ Nm. Cultures were diluted to an OD_720_ of 0.01−0.02 in the same medium, and 2.0 ml aliquots were placed in 12-well microtiter plates. Quadruplicate biological replicates of each culture were grown for up to 24 h at 30 °C with shaking and illumination at 50 to 80 μeinsteins m^−2^ s^−1^. The OD_720_ of each replicate was determined periodically during the 24–48 h growth period, and the change in OD_720_ versus time was used to determine generation times. Growth experiments were performed three times, and growth rates were expressed as the average and standard deviation for all replicates. Growth was exponential only up to an OD_720_ of about 0.15–0.20 and thereafter became linear because of light limitation due to self-shading by cells. Only exponential growth was used to determine the generation times.

### Plasmid construction

#### Cloning hsdRM and hsdS genes

The *hsdRM* and *hsdS* genes are located on plasmid D in *A. variabilis* (accession number NC_021575.1). A summary of plasmid constructions and primers is provided in Table 3. Plasmid pBP1207, which contained the *hsdRM* and *hsdS* genes with 1059 bp of upstream sequence in a pDU1-based plasmid, was constructed by joining adjacent fragments, which together comprised *hsdRM* and *hsdS* as well as 1059 bp upstream of *hsdRM*, into pUC18. One 2587-bp fragment containing a region that extended from 1059 bp upstream of *hsdRM* to the internal *hsdRM* BamH1 site was PCR amplified with primer pairs hsdRMS-P1L and hsdRMS-P1R. The second 3350-bp fragment, which extended from the BamHI site in the *hsdRM* gene to about 300 bp past the end of *hsdS*, was amplified using primer pairs hsdRMS-P2L and hsdRMS-P2R. The 5’ and 3’ ends of the 2587-bp upstream fragment were digested with HindIII and BamHI, respectively. The 5’ and 3’ ends of the downstream 3350-bp fragment were digested with BamHI and SacI, respectively. The two fragments, comprising the complete *hsdRMS*, were then ligated into the vector pUC18 (digested with HindIII and SacI) via a triple ligation to create plasmid pHC3. The 5.9-kb region of *hsdRMS* in pHC3 was sequenced to ensure that it had no mutations. The 5.9-kb HindIII/SacI fragment, containing the entire *hsdRMS* region, was excised from pHC3, blunted, and then cloned into the SmaI site of pRL57 to create pBP1207. Plasmids pBP1207 and control vectors lacking *hsdRMS* (pRL57 and pBP1310) were transformed into HB101 [pRL528] and then conjugated via triparental mating (Elhai et al., 1990) into cyanobacterial strains *Nostoc* sp. PCC 7120 and *Nostoc* sp. M131 to create strains *Nostoc* sp. PCC 7120 [pRL57], *Nostoc* sp. PCC 7120 [pBP1310], *Nostoc* sp. PCC 7120 [pBP1207], *Nostoc* sp. M131 [pRL57], *Nostoc* sp. M131 [pBP1310], and *Nostoc* sp. M131 [pBP1207].

**Table 3.**
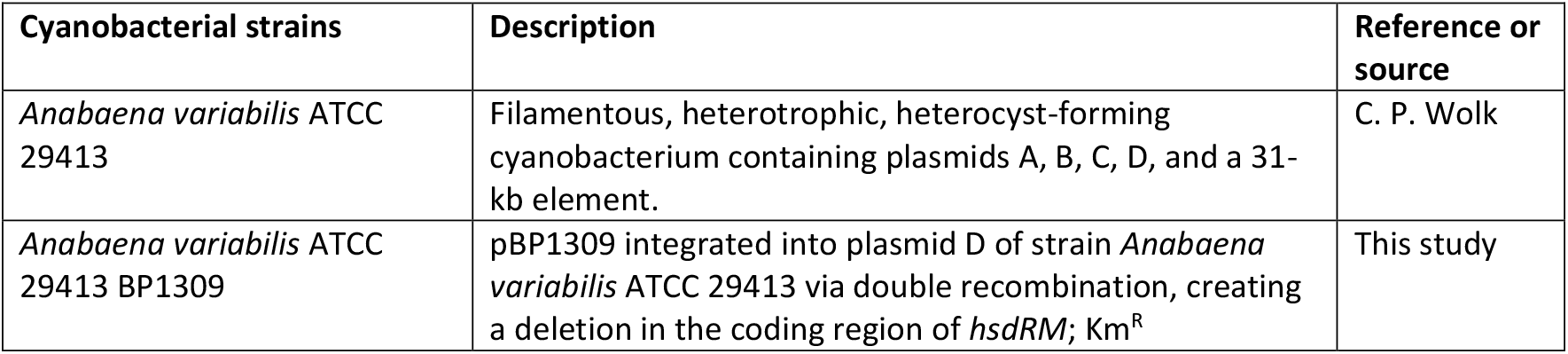
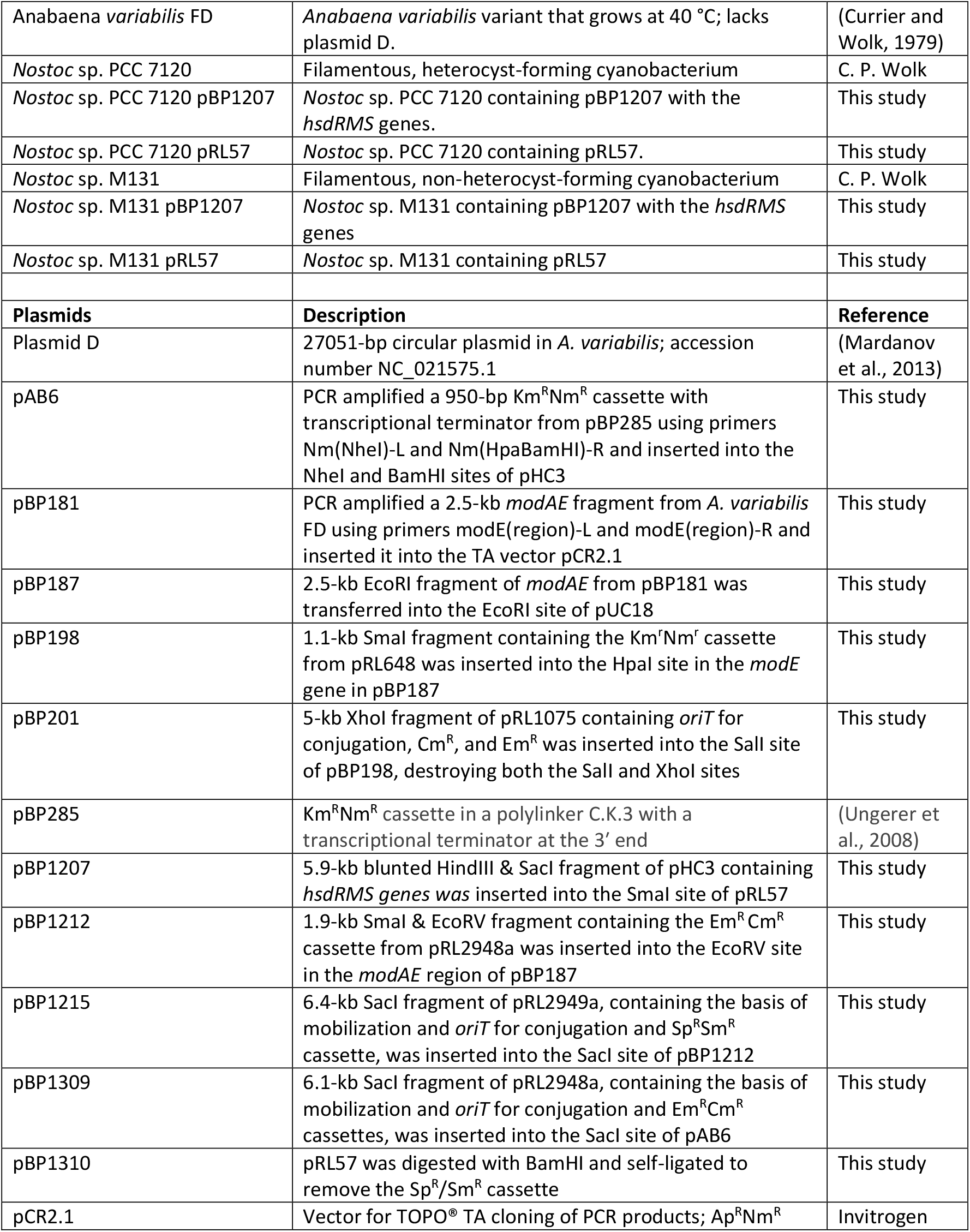
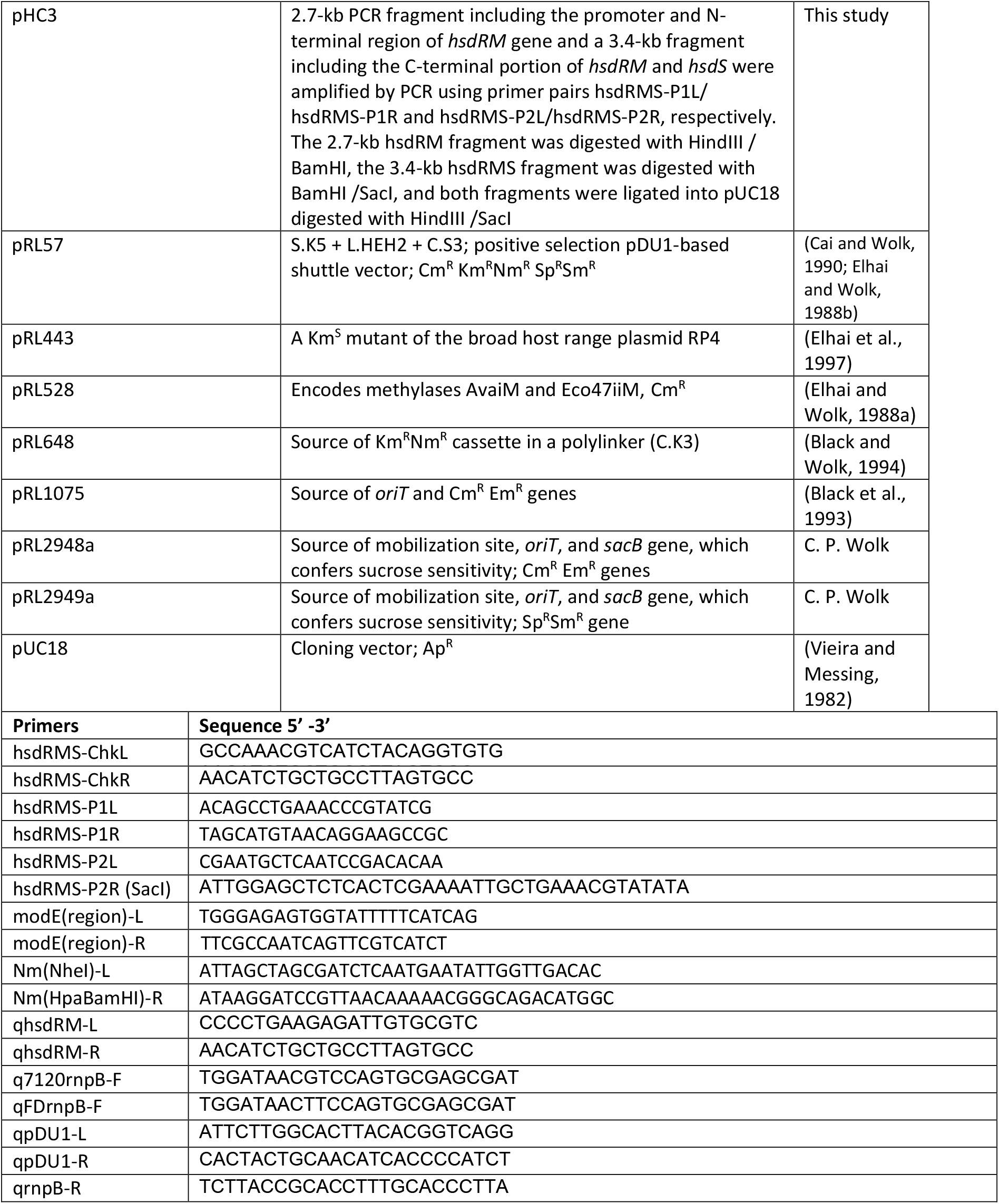
Strains, plasmids, and primers.

#### Construction of modAE plasmids for conjugation and recombination analysis

A 2.5-kb fragment from strain FD bearing *modE* (Ava2423) and *modA* (Ava2424) (molybdate transport genes) was PCR amplified using primers modE(region)-L and modE(region)-R and inserted into the TA vector pCR2.1 to create pBP181. The 2.5-kb EcoRI fragment from pBP181 containing the amplified *modAE* region was transferred to the vector pUC18, also digested with EcoRI, to create pBP187. A 1.1-kb SmaI fragment with the Nm^R^ cassette from pRL648 was inserted into the HpaI site within *modE* in pBP187 to create pBP198. Then, plasmid pBP201 was created by inserting a 5-kb XhoI fragment of pRL1075 (carrying *oriT*) into the SalI site of pBP198, destroying both the SalI and XhoI sites. Thus, the plasmid pBP201 has 1) the *modAE* region, interrupted by the Nm^R^ cassette—which, in previous experiments, has provided high-frequency recombination in the chromosome (Zahalak et al., 2004); 2) an *oriT*, required for transfer from *E. coli* to cyanobacteria by conjugation; and 3) an Em^R^ cassette in the vector to allow us to determine whether the conjugation event was by single recombination (producing Nm^R^ Em^R^ cells) or double recombination via loss of the vector portion (Nm^R^ Em^S^ cells). Thus, Nm^R^ colonies were tested for Em^R^ to quantify single-versus double-recombination events. A second plasmid, pBP1215, that differed in antibiotic resistance from pBP201 was also created—the Em^R^ Cm^R^ cassette was inserted in the *modAE* region, and the *aadA* Sp^R^Sm^R^ cassette was in the plasmid. First, pBP1212 was created by inserting a 1.9-kb SmaI/EcoRV Cm^R^ Em^R^ cassette from pRL2948a into the EcoRV site in the *modAE* region of pBP187 for selection of recombinants. Then pBP1215 was created by the addition of the 6.4-kb SacI fragment of pRL2949a, which contains *oriT* and the Sp^R^Sm^R^ cassette, into the SacI site of pBP1212. In constructing pBP1215, we overlooked the fact that pRL57 (the replicating control vector lacking *hsdRMS*) also carries Sp^R^Sm^R^. Thus, the mobilizable plasmid pBP1215 allowed us to identify single (Em^R^ Sp^R^Sm^R^) versus double recombinants (Em^R^ Sp^S^Sm^S^) in the *Nostoc* sp. PCC 7120 and *Nostoc* sp. M131 strains containing *hsdRM*. However, for the control strains with the plasmid but lacking *hsdRM*, we could not use the parent plasmid pRL57 because it confers Sp^R^Sm^R^, so double recombinants in the *modAE* region could not be identified, as the strain was always Sp^R^. Hence, we created the control plasmid pBP1310 by removing the Sp^R^Sm^R^ cassette from pRL57 by digesting pRL57 with BamHI and self-ligating the plasmid. This provided the control vector so that we could identify single versus double recombinants in the *modAE* region in *Nostoc* sp. PCC 7120 and *Nostoc* sp. M131.

#### Construction of hsdRM mutant

The 950-bp Km^R^Nm^R^ cassette containing a transcriptional terminator was amplified from pBP285 using primers Nm(NheI)-L and Nm(HpaBamHI) and inserted into the HpaI and BamHI sites in the *hsdRM* region of pHC3 from plasmid D in *Anabaena variabilis* to create pAB6. A 6.1-kb SacI fragment containing the mobilization site, *oriT, sacB*, and Em^R^ Cm^R^ cassettes from pRL2948 was inserted into the SacI site of pAB6 to create pBP1309. Plasmid pBP1309 was transformed into HB101 [pRL528] and then conjugated via triparental mating (Elhai et al., 1990) into cyanobacterial strain *Anabaena variabilis*. Recombinants were selected by Em^R^, and then verified to be double recombinant mutants by checking for Em^S^. Since the recipient strain was *A. variabilis*, all ex-conjugates were double-recombinant mutants. PCR using hsdRM-chkL/R primers was used to verify the deletion and that there were no wild-type copies of the region of interest. Cyanobacterial strain *A. variabilis* BP1309, lacking *hsdRM*, was conjugated via triparental mating with mobilizable plasmid pBP1215 carrying the *modAE* region interrupted by the Em^R^ cassette to determine the effect of the *hsdRM* region on conjugation and double recombination efficiency.

### Conjugation

*A. variabilis* and FD were conjugated with pBP201 using a triparental mating protocol. *Nostoc* sp. PCC 7120 and *Nostoc* sp. M131 strains containing either pBP1310 (pRL57 lacking Sp^r^Sm^r^ but containing *hsdRMS* genes) or pBP1207 (pRL57 containing *hsdRMS* genes), and *A. variabilis* BP1309 were conjugated with pBP1215 using a triparental mating protocol (Elhai et al., 1990). Cyanobacterial strains were grown to an OD_720_ of about 0.3 in 50 ml AA/8 media containing 5 mM NH_4_Cl, 10 mM TES, and 5 µg ml^-1^ Nm, when applicable, and *E. coli* cultures containing both the helper plasmid (HB101[pRL443]) and cargo plasmids (HB101[pBP201, pRL528] or HB101[pBP1215, pRL528]) were grown to log phase to an OD_600_ of about 0.5 in 25 ml L-broth lacking antibiotics. *Escherichia coli* and cyanobacterial cells were harvested, washed, and resuspended in 150 µl and 500 µl of AA/8, respectively. Aliquots of 75 µl of each *E. coli* strain (helper and cargo strains) and 150 µl of the cyanobacterial strain were combined, plated on a Millipore, 0.45 µM Triton-free MCE filter, and left for 72 h at 30°C with light. Cells were washed off the filter in AA/8 media, resuspended in 1 ml AA/8, and 100–300 µl volumes were plated and incubated on either BG-11 Nm^50^ plates (*A. variabilis* and FD) or BG-11 Nm^50^ Em^5^ plates (*Nostoc* sp. PCC 7120, *Nostoc* sp. M131, and *A. variabilis* BP1309) until colonies formed. Single (antibiotic-resistant) versus double (antibiotic-sensitive) recombination was determined by patching ex-conjugates with Em for *A. variabilis* and FD and SpSm for *Nostoc* sp. PCC 7120, *Nostoc* sp. M131, and *A. variabilis* BP1309 ex-conjugants.’

### RNA extraction and reverse transcriptase-qPCR (RT-qPCR)

*Nostoc* sp. PCC 7120 [pRL57], *Nostoc* sp. PCC 7120 [pBP1207], *Nostoc* sp. M131 [pRL57], and *Nostoc* sp. M131 [pBP1207] were grown in AA/8 media containing 5mM NH_4_Cl, 10 mM TES, pH 7.2, and 5 µg ml^-1^ neomycin. Cells were harvested by centrifugation, and RNA was isolated using the Tri Reagent (Sigma) as previously described (Ungerer et al., 2010). A total of 5 µg of RNA was subjected to DNase digestion (Turbo DNA-free kit; Ambion). A total of 40 ng of RNA was converted to cDNA in a 10 µl reaction mixture using iScript Reverse Transcription supermix for RT-qPCR (BioRad). The cDNA was diluted 1:20 to 0.2 ng µl^-1^. qPCR reactions were performed using 0.8 ng cDNA in a 10 µl reaction mixture with 5 pmol gene-specific primer and 1× SsoAdvanced SYBR Green supermix (BioRad). The following gene-specific primers were used: *rnpB*, qFDrnpB-F/qrnpB-R (*A. variabilis* and FD strains), q7120rnpB-F/qrnpB-R (7120 and M131 strains), and *hsdRM*, qhsdRM-L/R.

### Determination of loss of plasmids pRL57 and pBP1207

Triplicate cultures of *Nostoc* sp. PCC 7120 and *Nostoc* sp. M131 containing pRL57 (control) or pBP1207 (with *hsdRMS*) were grown in AA/8 medium containing 5 mM NH_4_Cl, 10 mM TES, pH 7.2, and 5 µg ml^-1^ Nm. Cells were washed to remove antibiotics and diluted in triplicate to an OD_720_ of 0.020 in AA/8 medium containing 5 mM NH_4_Cl and 10 mM TES, pH 7.2. Cultures were grown for a total of nine generations by diluting the cells 8-fold after every third generation, based on OD_720_. DNA was extracted as described above from cells harvested at 0 time and after nine generations of growth without Nm. PCR reactions were performed using 4 ng of chromosomal DNA in a 10 µl reaction mixture using 5 pmol of gene-specific primers and 1× SsoAdvanced SYBR Green Supermix (BioRad). The following primers were used: *rnpB*, q7120rnpB-F/qrnpB-R, and pDU1 plasmid, qpDU1-L/R.

### Determination of loss of plasmid D in *A. variabilis*

Cultures of *A. variabilis* grew in AA/8 medium containing 5 mM NH_4_Cl, 10 mM TES, pH 7.2 for over 30 generations at 40°C with light and shaking. After 30 generations, cultures were checked by qPCR for the ratio of plasmid D compared to chromosomal DNA using primers: *rnpB*, qFDrnpB-F/qrnpB-R, and *hsdRM*, qhsdRM-L/R. After no loss of plasmid D was observed *A. variabilis* filaments were sonicated down to 1–2 cell filaments, washed in AA/8, and then diluted in AA/8 medium containing 5 mM NH_4_Cl and 10 mM TES, pH 7.2, at dilutions ranging from 1/100 to 1/500,000. After a 1/100,000 dilution, when the culture reached an OD_720_ of∼0.1, it was sonicated again to short filaments, the cells were washed and plated on BG-11 plates at various dilutions for single colonies. We picked 50 of the smallest colonies obtained from the sonication for semi-quantitative colony PCR checking for the presence of plasmid D, using qFDrnpB-F/R and hsdRMS-ChkL/R primers. All colonies had a PCR product using *hsdRMS* primers; we picked two colonies that had the lightest bands and did qPCR on those colonies using *rnpB*, qrnpB-F/qrnpB-R, and *hsdRM*, qhsdRM-L/R primers.

## Data availability

All data used in the experiments are provided here.

## Acknowledgment

This work was supported by the National Science Foundation grant MCB-1818298.

